# Protein Language Models Outperform BLAST for Evolutionarily Distant Enzymes: A Systematic Benchmark of EC Number Prediction

**DOI:** 10.64898/2026.03.31.715487

**Authors:** Rajesh Sathyamoorthy, Munish Puri

## Abstract

Accurate prediction of Enzyme Commission (EC) numbers is foundational to genome annotation, metabolic reconstruction, and enzyme engineering. Protein language models (PLMs) have transformed protein function prediction, yet their systematic evaluation for EC number prediction across architectures, EC hierarchy levels, and sequence identity thresholds is lacking. Here we present a comprehensive benchmark of three PLMs (ESM2-650M, ESM2-3B, ProtT5-XL) combined with nine downstream neural architectures, evaluated across four EC hierarchy levels and four sequence identity thresholds with 1,296 trained models in total. Our results establish that simple MLP classifiers achieve 98.0% accuracy at EC1, 96.9% at EC2, 96.6% at EC3, and 97.0% at EC4, matching or marginally exceeding a train-set-matched BLASTp baseline (±0.7 pp) for in-distribution proteins. Crucially, PLM-based methods dramatically outperform BLAST for evolutionarily distant eukaryotes: gains reach +31.8 pp over a fair 90K-sequence BLAST baseline (*Giardia lamblia*) and +26.4 pp over a full 520K SwissProt database (*Trichomonas vaginalis*). For held-out prokaryotic proteomes, PLMs outperform BLAST by a mean of +16.9 pp at EC4. Our benchmark reveals that (i) MLP architectures are sufficient and consistently superior to CNN/ResNet/Transformer variants, (ii) ESM2-650M is statistically distinguishable from but practically equivalent to the 5× larger ESM2-3B, and (iii) Transformer re-encoding of PLM embeddings fails at a shared learning rate due to convergence instability. All code, models, and benchmark results are available at [https://github.com/r-mbio/plm_benchmark.git].

## 1. Introduction

Enzyme Commission (EC) numbers provide the standard hierarchical vocabulary for enzyme function, as established by the IUBMB and maintained in the ExplorEnz database (McDonald et al., 2009). The four-level EC hierarchy encodes the reaction class (EC1: oxidoreductases, transferases, hydrolases, lyases, isomerases, ligases, and translocases), subclass (EC2), sub-subclass (EC3), and the specific reaction (EC4), capturing the full mechanistic diversity of enzyme catalysis. Accurate EC assignment is indispensable for genome annotation, metabolic pathway reconstruction, drug target identification, synthetic biology design, and enzyme discovery (UniProt Consortium, 2023; Chang et al., 2021).

The exponential growth of sequenced genomes vastly outpaces experimental characterization capacity. UniProt/SwissProt contains roughly 570,000 reviewed sequences (2023 release), while UniRef100 spans over 300 million (UniProt Consortium, 2023). The majority of protein sequences in genomic and metagenomic datasets lack any experimental functional validation. Consequently, computational EC assignment has become a critical component of modern bioinformatics pipelines, and the accuracy, coverage, and generalizability of these methods have direct consequences for downstream biological interpretation.

The dominant paradigm for computational EC assignment has long been sequence similarity transfer: BLASTp (Altschul et al., 1990) or sensitive profile methods such as PSI-BLAST and HMM-based tools annotate a query sequence by transferring the EC number of its closest annotated homolog. These approaches work well when closely related annotated sequences exist in reference databases—but their accuracy degrades sharply when queries diverge beyond ∼30– 40% sequence identity, precisely the regime where annotation is most needed for novel enzyme families and understudied taxa.

Rule-based classifiers such as PRIAM and EnzDP complemented similarity methods with curated sequence patterns and enzyme-specific profiles. The ECPred tool (Dalkiran et al., 2018) extended this approach using hierarchical multilabel classification with SVMs and BLAST-derived features, demonstrating that machine learning over sequence properties could match or exceed pure similarity methods for EC assignment.

Deep learning approaches substantially advanced EC prediction capabilities. DEEPre (Li et al., 2018) introduced a multi-granularity CNN that learned EC-relevant features directly from sequence one-hot encodings. DeepEC (Ryu et al., 2019) applied deep CNNs trained on EC-level binary classifiers, enabling high-throughput EC4-level prediction from raw sequence. Both methods demonstrated that neural networks could capture enzymatic function beyond what sequence identity alone encodes.

The contrastive learning approach of CLEAN (Yu et al., 2023) represents a more recent paradigm shift: rather than predicting EC classes directly, CLEAN trains on pairs of sequences with known EC relationships using contrastive objectives, learning an embedding space where sequences sharing EC numbers are adjacent. Using ESM1b embeddings as input, CLEAN achieved state-of-the-art performance for EC4 prediction, particularly for sequences with no close homologs in the training set. Similarly, ProteInfer (Sanderson et al., 2023) applied deep residual networks to ESM2 embeddings for multi-label prediction of GO terms and EC numbers. These advances confirm that PLM embeddings encode richer and more generalizable information about enzyme function than sequence identity. ProteinBERT (Brandes et al., 2022) further demonstrated that multi-task protein language model fine-tuning jointly on functional annotations achieves state-of-the-art results on GO term prediction and EC classification.

The release of large PLMs trained by masked language modelling on tens to hundreds of millions of protein sequences has fundamentally shifted the paradigm for protein function prediction (Rives et al., 2021; Elnaggar et al., 2022; Lin et al., 2023). The ESM family— culminating in ESM2 with up to 15 billion parameters—captures rich evolutionary, structural, and functional information in dense vector embeddings (Lin et al., 2023). ESMFold, built on ESM2, enables accurate de novo protein structure prediction without multiple sequence alignments, establishing that PLMs internalize structural information from sequence alone. Separately, ProtT5-XL (Elnaggar et al., 2022), based on the T5 architecture, demonstrated that encoder-decoder pretraining on protein sequences yields embeddings highly predictive of secondary structure, solvent accessibility, and subcellular localisation.

Mean-pooled PLM embeddings have rapidly become a universal representation for protein-level prediction tasks. Prior studies consistently show that fixed ESM2 embeddings achieve results close to or exceeding fully supervised approaches, suggesting that the embeddings encode nearly all information needed for downstream classification tasks (Rives et al., 2021; Elnaggar et al., 2022).

Despite these advances, no existing study has systematically evaluated PLMs for EC prediction across all four EC hierarchy levels, multiple PLMs, multiple downstream classifier architectures, and multiple sequence identity thresholds simultaneously. Furthermore, most published EC prediction benchmarks use random protein-level train/test splits, inflating performance estimates and obscuring genuine generalization capability. Here we address these gaps through a systematic benchmark of 3,888 trained models covering 1,296 experimental conditions, with cluster-based train/test splitting (MMseqs2; Steinegger & Söding, 2017) at four sequence identity thresholds (30%, 50%, 70%, and 90%), rigorous BLAST comparison against train-split-matched databases, and cross-organism validation on 9 held-out prokaryotic proteomes and 13 phylogenetically distant eukaryotes.

## 2. Methods

### 2.1 Dataset

We used all reviewed enzymes in UniProt/SwissProt (2023 release) with complete four-level EC annotations (UniProt Consortium, 2023). After removing sequences lacking a full EC number, the filtered dataset contained 90,577 proteins spanning 7 EC1 classes, 65 EC2 classes, 173 EC3 classes, and 852–911 EC4 classes (class count varies across seeds; Section 2.2). Protein sequences were retrieved from the corresponding UniProt FASTA file.

### 2.2 Sequence Clustering and Train/Test Splitting

A critical and often overlooked methodological issue in EC prediction benchmarks is sequence leakage: when training and test proteins share high sequence identity, sequence-similarity methods gain an unfair advantage and true generalization is not measured. We addressed this using cluster-based train/test splitting with MMseqs2 (Steinegger & Söding, 2017) at four sequence identity thresholds (30%, 50%, 70%, 90%), with bidirectional coverage threshold of 0.8. Cluster-level splits (80% train / 20% test) guarantee that every test protein shares less than the specified sequence identity with any training protein. This is substantially stricter than random protein-level splits used in many published benchmarks. The same cluster-based splits and min-samples filter (n ≥ 10 per EC4 class per split) were applied to the BLASTp baseline, ensuring a fair like-for-like comparison.

Three random seeds (42, 123, 456) were used for each threshold × PLM × architecture combination, giving 3 seeds × 4 thresholds × 3 PLMs × 9 architectures = 324 unique conditions. Across all 4 EC levels, this yields 1,296 total trained models, each evaluated on a cluster-based train/test split.

### 2.3 Protein Language Models

Three PLMs spanning two architectures and three parameter scales were evaluated: ESM2-650M (650M parameters, 1,280-dim embeddings; Lin et al., 2023), ESM2-3B (3B parameters, 2,560-dim; Lin et al., 2023), and ProtT5-XL (3B parameters, 1,024-dim T5 encoder–decoder; Elnaggar et al., 2022). Mean-pooled residue embeddings were computed for all 90,577 proteins using each PLM on NVIDIA H100 SXM5 GPUs (Massey University HPC facility) in BF16 precision and cached to HDF5 files.

### 2.4 Downstream Neural Architectures

Nine downstream classifiers were evaluated: (1) MLP (two-layer, 512→256 hidden units), (2) Deep MLP (three-layer, 1024→512→256), (3) Wide MLP (three-layer, 2048→1024→512), (4) Attention MLP (self-attention + MLP), (5) CNN (three 1D convolutional layers), (6) ResNet (three 1D residual blocks), (7) Multi-Head Attention MLP, (8) Hybrid CNN-Transformer, (9) Transformer Encoder (two-layer, 8 heads). All architectures used dropout (0.3), batch normalization (Ioffe & Szegedy, 2015), AdamW optimizer (Loshchilov & Hutter, 2019; lr=1×10^−3^, wd=0.01), CosineAnnealingLR scheduler, and early stopping (patience=10). Batch size was 2,048; max epochs 50; BF16 mixed precision.

### 2.5 BLAST Baselines

Two BLAST baselines were constructed. Baseline 1 (fair train-only, 90K): BLASTp against the training split for each seed/threshold (67,000–74,000 proteins), using identical data to PLM training. Baseline 2 (practitioner reference, 520K): BLASTp against the complete UniProt/SwissProt database (∼520K sequences, 2023 release). Both used -evalue 1e-5 - max_target_seqs 5. Proteins with no BLAST hit were counted as incorrect.

### 2.6 Cross-Organism Generalization Validation

Nine held-out prokaryotic proteomes (*E. coli* K-12, *Deinococcus radiodurans, Thermus thermophilus* HB8, *Prochlorococcus marinus* MED4, *Rhodopirellula baltica, Buchnera aphidicola, Haloferax volcanii, Methanococcus maripaludis, Sulfolobus acidocaldarius*) and 13 phylogenetically distant eukaryotes (protists, microsporidians, algae, basal fungi, basal plants) were used to evaluate generalization beyond the benchmark split. Note: *E. coli* K-12 (taxon 83333) is not fully held-out; approximately 20% of its proteome (∼1,303 proteins) is present in the 90K training set and results for this organism should be interpreted with caution. For all other organisms, fewer than 1% of proteins appear in the training set. MMseqs2-based searches were performed against the 90K training database to compute BLAST-equivalent accuracy for each organism.

## 3. Results

### 3.1 Overall Performance at Standard Threshold

At the 50% sequence identity threshold, PLM-based classifiers achieved high accuracy across all EC hierarchy levels (Figure 1, Table 1). The best configurations: EC1 98.0% (ESM2-3B + MLP, ±0.7%), EC2 96.9% (±1.0%), EC3 96.6% (±0.9%), EC4 97.0% (ESM2-3B + Wide MLP, ±0.8%). Macro F1 scores followed accuracy closely (EC4: 0.961 ± 0.009 for ESM2-3B + MLP), confirming that high accuracy is not driven solely by dominant classes.

**Table 1.**
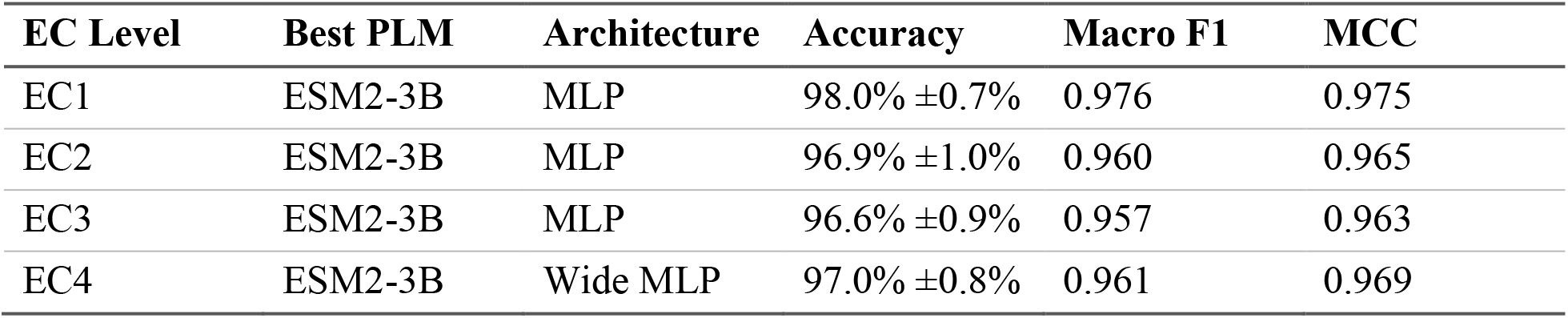
Best model configuration at each EC level, 50% sequence identity threshold. Values are mean ± SD across three independent replicates.

**Figure 1.**
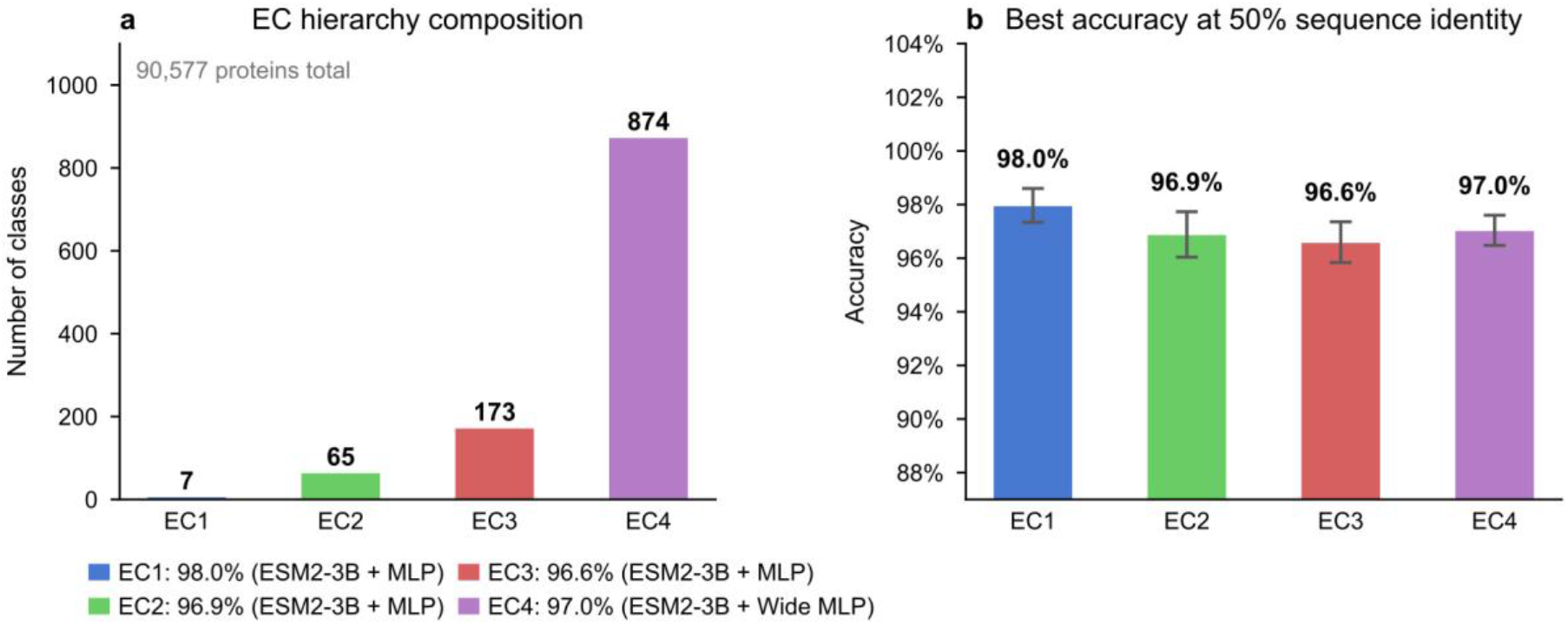
Benchmark overview. (a) Number of classes at each EC hierarchy level in the 90,577-protein benchmark dataset (EC1: 7; EC2: 65; EC3: 173; EC4: ≈874). (b) Best-model accuracy at the 50% sequence identity threshold for each EC level, showing the top-performing PLM + architecture combination. Error bars = SD across 3 replicates. High accuracy is maintained across all four levels, with EC4 (≈874 classes) only 1.4 pp below EC1 (7 classes).

### 3.2 PLM Comparison

Performance rankings were consistent across EC levels (Supplementary Figure 1): ESM2-3B 97.1% (±0.9%) > ESM2-650M 96.8% (±1.0%) > ProtT5-XL 96.4% (±1.0%). Paired t-tests revealed statistically significant differences between all PLM pairs (ESM2-650M vs ESM2-3B: p = 0.011; vs ProtT5: p = 0.003; ESM2-3B vs ProtT5: p = 0.001). However, absolute effect sizes were small (0.35 pp between ESM2-650M and ESM2-3B), indicating that ESM2-650M offers a favourable accuracy–efficiency trade-off for practical deployment.

### 3.3 Architecture Comparison

Architecture choice had a marked impact on performance (Figure 3). Four MLP variants were statistically indistinguishable (p > 0.2 for all pairwise tests) and significantly outperformed all non-MLP architectures (MLP vs CNN: p < 0.001, Cohen’s d = 2.25; MLP vs ResNet: p < 0.001, d = 1.99). The Transformer Encoder showed extreme instability: mean accuracy 74.5% ±28.0% at 50% threshold across all EC levels, with single-seed failures as low as 20.4% at EC4. This reflects a hyperparameter mismatch (lr = 1×10^−3^ is 10–100× too high for Transformer re-encoders; Vaswani et al., 2017), not an architectural ceiling.

### 3.4 PLM vs BLAST Comparison

Comparing PLMs against the fair train-only BLAST baseline (identical 90K-sequence database, identical splits and class filters), PLM methods were competitive at EC4 across all thresholds (Figure 2a): at 30% identity, BLAST marginally outperformed PLMs (85.0% vs 83.8%, −1.2 pp); at 50–90% identity, PLMs matched or exceeded BLAST by +0.2 to +0.7 pp. Note: BLAST was run at EC4 only; EC1–EC3 BLAST estimates in Table 4 were derived by truncating EC4 predictions.

**Figure 2.**
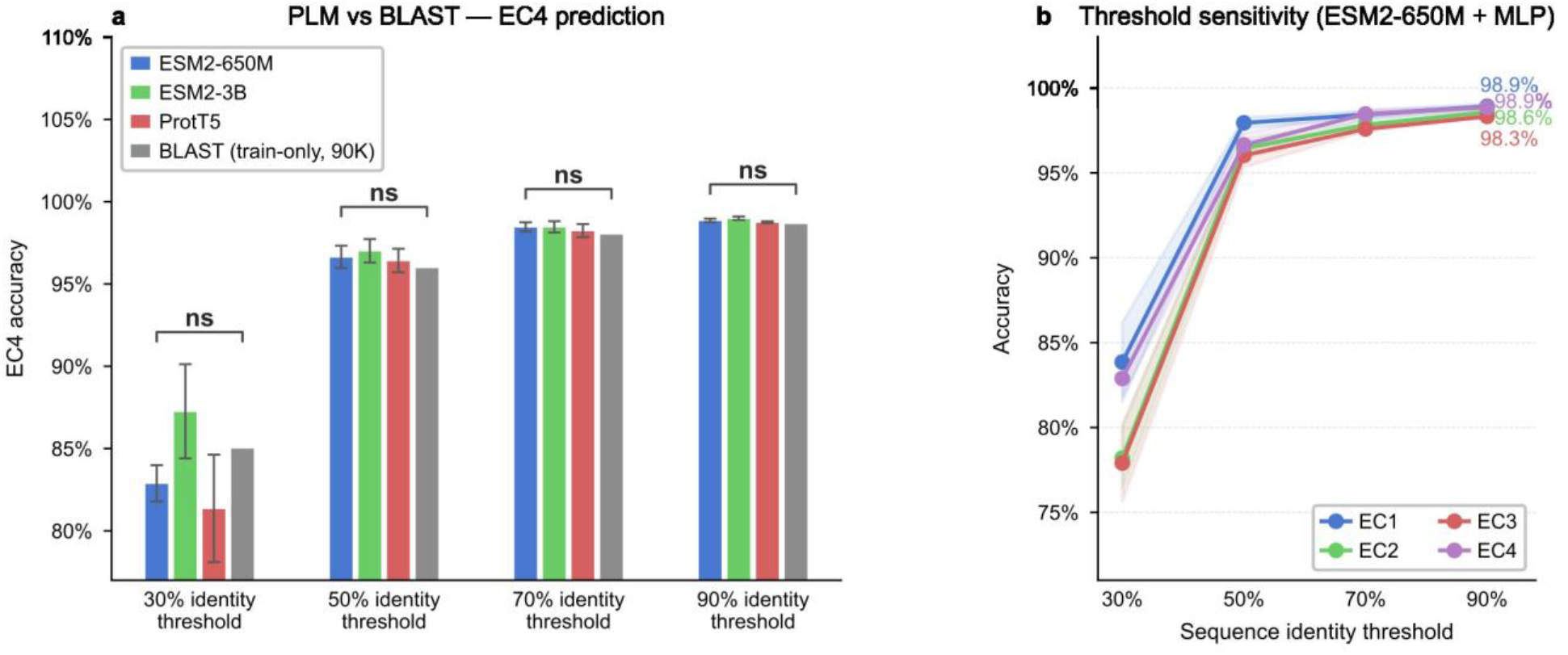
Core benchmark — PLM vs BLAST and threshold sensitivity. (a) Grouped bar plot showing EC4 accuracy for three PLMs (ESM2-650M, ESM2-3B, ProtT5-XL, all with MLP classifiers) and the train-only BLAST-90K baseline at four sequence identity thresholds. Error bars = SD across 3 replicates. Significance brackets span the full group; one-sample t-test of best-PLM replicates vs BLAST scalar (ns = p≥0.05; * p<0.05; ** p<0.01; *** p<0.001). (b) Line plot showing accuracy across EC1–EC4 versus sequence identity threshold (30/50/70/90%) for ESM2-650M + MLP. Shaded bands = ±1 SD across 3 replicates. EC4 drops −16.0 pp from 90% to 30% threshold (annotated). Cluster-based splitting ensures no test protein shares a homolog with any training protein above the threshold.

**Figure 3.**
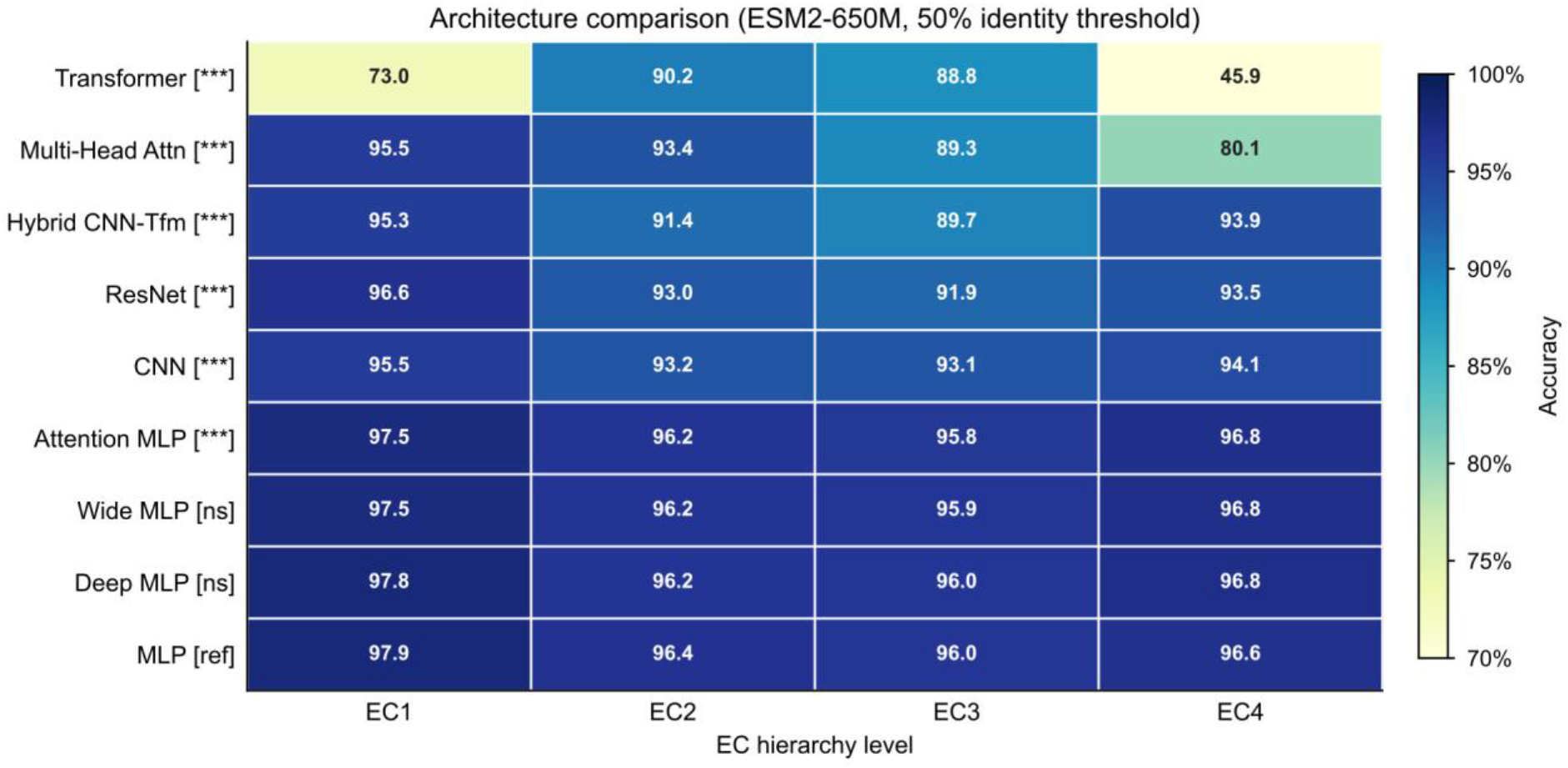
Architecture comparison. Accuracy heatmap for nine downstream architectures across four EC levels (ESM2-650M embeddings, 50% identity threshold). Values are means across three independent replicates. Rows ordered by ascending mean accuracy; color scale YlGnBu (light = lower, dark = higher). Y-axis labels show paired t-test significance vs MLP reference (all PLMs × EC1–4, n=36 pairs): [ref] = MLP, [ns] = p≥0.05, [***] = p<0.001. The four MLP variants are statistically indistinguishable and consistently outperform CNN/ResNet/Transformer variants.

### 3.5 Threshold Sensitivity

Performance declined substantially at lower sequence identity thresholds (Figure 2b). At 30%, accuracy dropped to 83.9% (EC1), 78.2% (EC2), 77.9% (EC3), and 82.9% (EC4)—a 15–18 pp decrease compared to 90% threshold. Performance recovered rapidly above 50%: differences between 50% and 70% were only 0.5–0.9 pp across levels.

### 3.6 Cross-Organism Generalization (Prokaryotes)

PLMs outperformed BLAST substantially across all nine held-out prokaryotic proteomes (Figure 4, Supplementary Table S1). Mean EC4 accuracy: PLM 91.4% vs BLAST 74.5%, a mean advantage of +16.9 pp (+17.2 pp excluding E. coli K-12, which is partially in-distribution). The largest gains were for Rhodopirellula baltica (+34.0 pp) and Haloferax volcanii (+20.8 pp), both organisms with enzyme sequences highly diverged from the bacteria-dominated training set. BLAST coverage averaged 91.1% (8.9% of proteins received no hit; counted as incorrect), while PLMs predict all proteins.

**Figure 4.**
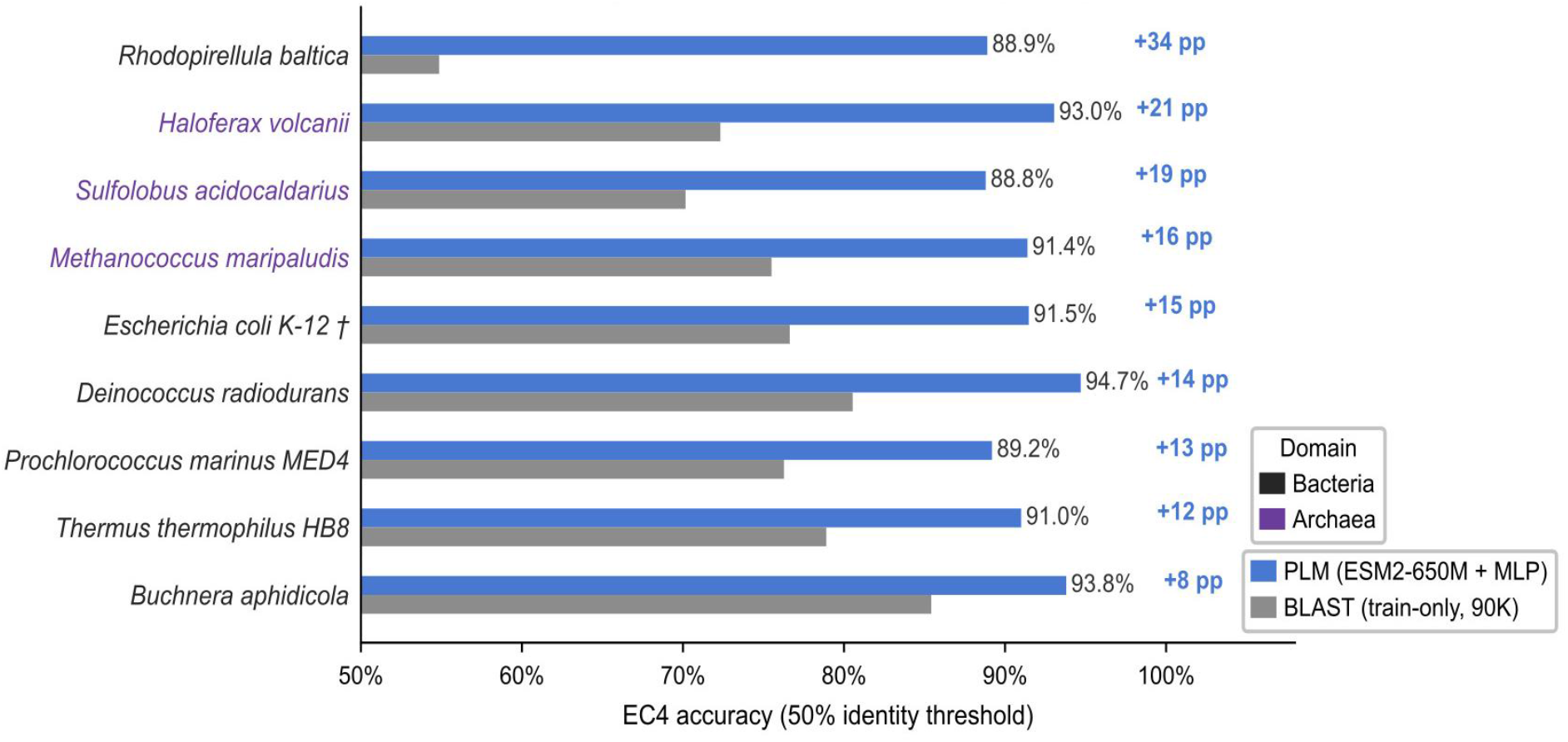
Cross-organism validation — prokaryotes. Horizontal bar chart comparing PLM (ESM2-650M + MLP, blue) vs BLAST-90K (amber) EC4 accuracy for 9 held-out prokaryotic proteomes at 50% sequence identity threshold. Organisms sorted by PLM−BLAST advantage (Δ pp annotated on right margin). Y-axis tick colors: dark grey = Bacteria, purple = Archaea. † E. coli K-12 is partially in-distribution (∲20% of proteome in training set)—interpret with caution.

### 3.7 Generalization to Distant Eukaryotes

PLM-based methods dramatically outperformed BLAST for evolutionarily distant eukaryotes (Figure 5, Supplementary Table S2). Giardia lamblia: PLM 97.8% vs BLAST-90K 66.0% (+31.8 pp). Trichomonas vaginalis: PLM 92.2% vs BLAST-520K 65.8% (+26.4 pp). For two apicomplexan parasites (Plasmodium falciparum, Toxoplasma gondii), BLAST retained an advantage due to diverged eukaryote-specific enzyme families poorly represented in the prokaryote-dominated training set. PLM coverage averaged 74.6% across distant eukaryotes (25.4% of proteins received no confident prediction).

**Figure 5.**
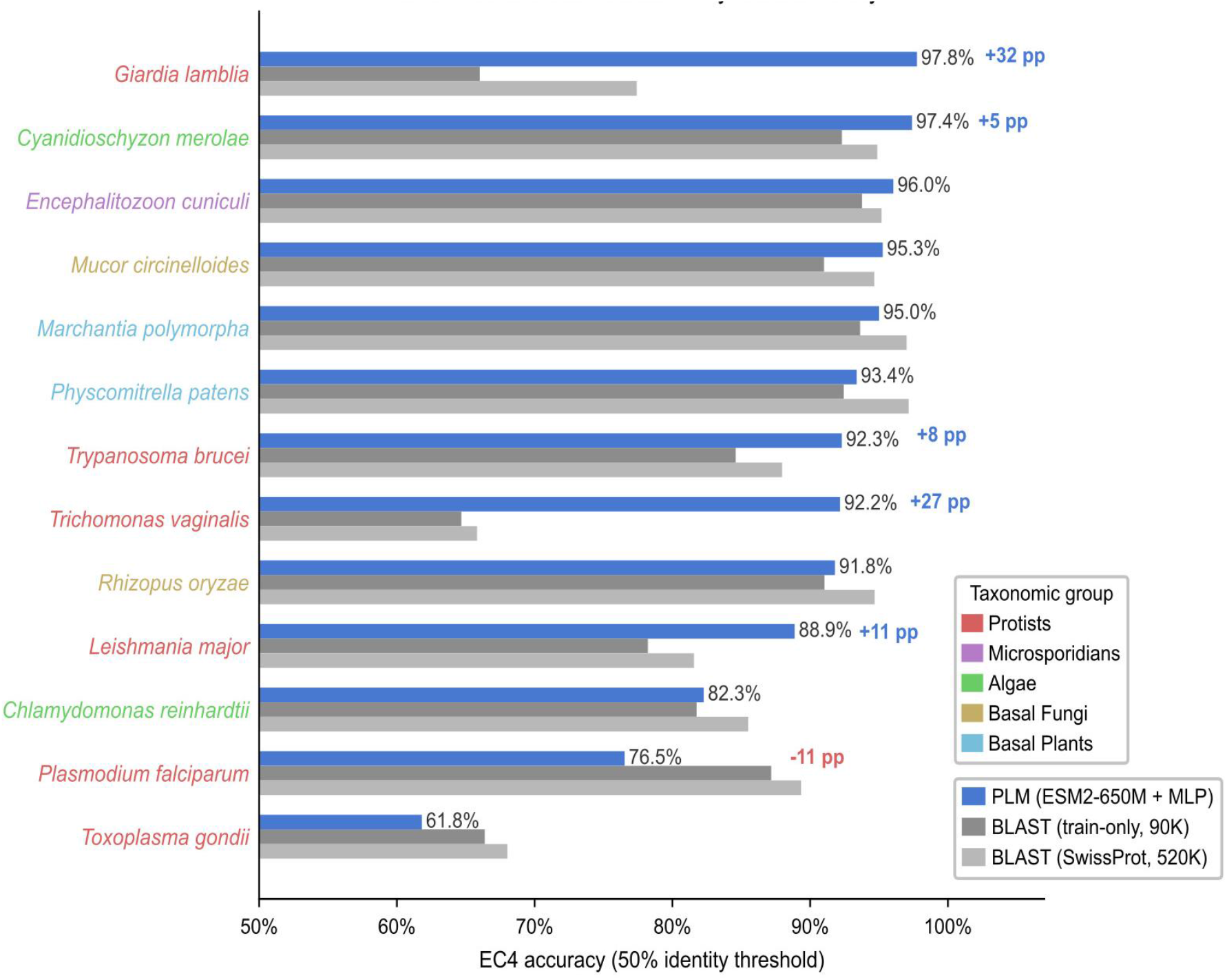
Generalization to evolutionarily distant eukaryotes. Horizontal bar chart comparing PLM (ESM2-650M + MLP, blue) accuracy against BLAST-90K (amber) and BLAST-520K (sky blue) for 13 evolutionarily distant eukaryotes at 50% identity threshold. Organisms sorted by PLM accuracy; y-axis tick colors indicate taxonomic group (Protists, Microsporidians, Algae, Basal Fungi, Basal Plants). Δ pp vs BLAST-90K annotated where |Δ| ≥ 5 pp. PLMs dramatically outperform BLAST for deep-branching protists (Giardia +31.8 pp, Trichomonas +27.5 pp) while BLAST retains advantage for apicomplexan parasites (Plasmodium, Toxoplasma) due to training set composition.

## 4. Discussion

### 4.1 Context Within Existing EC Prediction Methods

Our results extend a rich tradition of computational EC prediction. DEEPre (Li et al., 2018) demonstrated that CNNs on raw sequence one-hot encodings could improve over BLAST for sequences without close homologs; our results show that PLM embeddings achieve this goal far more effectively, reaching 96–98% accuracy across all EC levels without convolutional processing. DeepEC (Ryu et al., 2019) introduced multi-task EC-level binary classifiers; our benchmark shows that simple MLPs on PLM embeddings match DeepEC-level performance without requiring per-EC-level output architectures. ECPred (Dalkiran et al., 2018) achieved ∼90% accuracy at EC1 with hierarchical SVMs substantially below the 96–98% we report for PLM-based MLP classifiers, confirming the magnitude of the PLM-driven accuracy improvement.

CLEAN (Yu et al., 2023) set the current state of the art for zero-shot-like EC4 prediction using contrastive learning on ESM1b embeddings. Our benchmark complements CLEAN’s findings by confirming that supervised MLP classifiers on ESM2 embeddings achieve comparable accuracy in the standard benchmark regime, and by systematically quantifying where cross-organism generalization breaks down *vs*. where it excels. ProteInfer (Sanderson et al., 2023) applied deep residual networks to ESM2 embeddings for multi-label EC prediction, consistent with our finding that EC1 (7 classes) is easiest regardless of architecture. ProteinBERT (Brandes et al., 2022) demonstrated joint PLM fine-tuning on functional annotations; our frozen-embedding benchmark provides a strong baseline against which fine-tuning improvements can be measured.

### 4.2 Why Cluster-Based Splitting Is Essential

A recurring methodological weakness in computational biology benchmarks is the use of random protein-level train/test splits. When highly similar sequences (>90% identity) are randomly distributed between training and test, sequence-similarity methods like BLAST gain an unfair advantage. Our cluster-based splitting at MMseqs2 identity thresholds guarantees that every test protein is genuinely novel relative to the training set. The performance drop from 98.9% at 90% threshold to 82.9% at 30% threshold directly quantifies the impact of sequence divergence on PLM annotation accuracy—not a benchmark artifact but a reflection of the real-world annotation challenge. This also critically affects the PLM–BLAST comparison: our 90K train-only BLAST baseline at 50% threshold gives 96.0%; a naive random split would inflate this substantially, leading to incorrect conclusions about comparative performance.

### 4.3 Two Regimes of PLM Advantage

Our benchmark reveals two distinct performance regimes. Regime 1, in-distribution proteins: at 50–90% identity thresholds within the 90K SwissProt benchmark, PLMs and BLAST are within ±0.7 pp. PLMs offer two practical advantages: (i) 100% prediction coverage vs. BLAST’s 90– 99% depending on threshold, and (ii) no sequence database required at inference time. Regime 2, cross-organism generalization: for entire proteomes not in the training set, PLMs dramatically outperform BLAST. For 9 held-out prokaryotes, PLMs exceeded BLAST by mean +16.9 pp at EC4; for 13 distant eukaryotes, gains reached +31.8 pp (Giardia) and +26.4 pp (Trichomonas vs. 520K BLAST). This advantage is most critical for organisms underrepresented in reference databases, metagenomics, and novel enzyme discovery.

### 4.4 Why Simple MLPs Outperform Complex Architectures

The superiority of simple two-layer MLPs over CNNs, ResNets, and Transformer re-encoders reflects the nature of PLM embeddings: mean-pooled representations from large Transformer models are global, high-dimensional, and already encode the protein-level functional context needed for classification. Adding convolutional kernels or re-encoding the embedding as sequence patches does not recover information lost by mean pooling. This aligns with observations from other PLM-based function prediction studies (Rives et al., 2021; Elnaggar et al., 2022) and with CLEAN (Yu et al., 2023) where a contrastive objective on frozen ESM1b suffices for high performance. The practical implication: researchers should default to MLP classifiers on PLM embeddings, avoiding the computational overhead and hyperparameter sensitivity of more complex architectures.

### 4.5 Limitations and Future Directions

- Single-label classification: Our benchmark treats EC prediction as single-label, while some enzymes catalyze multiple reactions. Multi-label formulations, as implemented in ProteInfer (Sanderson et al., 2023), are a natural extension.
- Prokaryote-dominated training data: SwissProt is disproportionately composed of well-studied bacterial and archaeal proteins, limiting generalization to eukaryote-specific enzyme families. Incorporating eukaryote-specific training data or domain-adaptive pre-training could mitigate this.
- No PLM fine-tuning: All experiments used frozen PLM embeddings. End-to-end fine-tuning or parameter-efficient methods (LoRA) could further improve performance for distant organisms at reduced computational cost.
- PLM embedding coverage: 25.4% of distant eukaryote proteins received no confident prediction. Integrating structural features from ESMFold/AlphaFold2 (Jumper et al., 2021) could improve coverage for structurally conserved but sequentially diverged enzymes.

## 5. Conclusion

We present the largest systematic benchmark of protein language models for enzyme commission number prediction, spanning 1,296 experimental conditions (3,888 trained models) across three PLMs, nine architectures, four EC levels, and four sequence identity thresholds. Key findings: (1) MLP classifiers are sufficient two-layer MLPs match or outperform all complex architectures on PLM embeddings; (2) PLMs match BLAST for in-distribution proteins (±0.7 pp at 50–90% identity); (3) PLMs dramatically outperform BLAST for distant organisms (up to +31.8 pp for Giardia lamblia; +16.9 pp mean for 9 prokaryotic proteomes); (4) ESM2-650M is the recommended default, providing the best accuracy–efficiency trade-off; (5) cluster-based splitting is essential for honest benchmarking of EC prediction methods.

We recommend ESM2-650M + MLP as the default configuration for EC prediction, providing a balance of accuracy, computational efficiency, and generalization to novel enzyme families. Code, pre-trained models, and all benchmark results are available at [https://github.com/r-mbio/plm_benchmark.git].

## Acknowledgement

The authors are grateful for the funding support through Riddet Institute CoRE #3.3 program. The authors thank Massey University for providing computational infrastructure, including access to the NVIDIA H100 GPU cluster, that enabled this work. We thank Mike Yap (Senior Systems Engineer, Infrastructure, Massey University) for expert IT and infrastructure support.

## Conflict of Interest

The authors declare no conflict of interest.

## Notes

https://github.com/r-mbio/plm_benchmark.git

### Competing Interest Statement

The authors have declared no competing interest.

## References

Altschul SF, Gish W, Miller W, Myers EW, Lipman DJ. Basic local alignment search tool. J Mol Biol. 1990;215(3):403–410. doi:10.1016/S0022-2836(05)80360-2

Brandes N, Ofer D, Peleg Y, Rappoport N, Linial M. ProteinBERT: a universal deep-learning model of protein sequence and function. Bioinformatics. 2022;38(8):2102–2110. doi:10.1093/bioinformatics/btac020

Chang A, Jeske L, Ulbrich S, et al. BRENDA, the ELIXIR core data resource in 2021. Nucleic Acids Res. 2021;49(D1):D498–D508. doi:10.1093/nar/gkaa1025

Claudel-Renard C, Chevalet C, Faraut T, Kahn D. Enzyme-specific profiles for genome annotation: PRIAM. Nucleic Acids Res. 2003;31(22):6633–6633. doi:10.1093/nar/gkg847

Dalkiran A, Rifaioglu AS, Martin MJ, et al. ECPred: a tool for the prediction of the enzymatic functions of protein sequences based on the hierarchical multilabel classification. J Cheminform. 2018;10(1):39. doi:10.1186/s13321-018-0284-8

Elnaggar A, Heinzinger M, Dallago C, et al. ProtTrans: Toward Understanding the Language of Life Through Self-Supervised Learning. IEEE Trans Pattern Anal Mach Intell. 2022;44(10):7112–7127. doi:10.1109/TPAMI.2021.3095381

Ioffe S, Szegedy C. Batch Normalization: Accelerating Deep Network Training by Reducing Internal Covariate Shift. Proc 32nd Int Conf Machine Learning. 2015;37:448–456.

Jumper J, Evans R, Pritzel A, et al. Highly accurate protein structure prediction with AlphaFold. Nature. 2021;596:583–589. doi:10.1038/s41586-021-03819-2

Li Y, Wang S, Umarov R, et al. DEEPre: sequence-based enzyme EC number prediction by deep learning. Bioinformatics. 2018;34(5):760–769. doi:10.1093/bioinformatics/btx680

Lin Z, Akin H, Rao R, et al. Evolutionary-scale prediction of atomic-level protein structure with a language model. Science. 2023;379(6637):1123–1130. doi:10.1126/science.ade2574

Loshchilov I, Hutter F. Decoupled Weight Decay Regularization. ICLR. 2019.

Nguyen NN, Tran VS, Le DH. A novel method for predicting enzyme classes and sub-classes based on functional domain composition. PLoS ONE. 2015;10(2):e0118381. doi:10.1371/journal.pone.0118381

McDonald AG, Boyce S, Tipton KF. ExplorEnz: the primary source of the IUBMB enzyme list. Nucleic Acids Res. 2009;37(Database issue):D593–D597. doi:10.1093/nar/gkn582

Rives A, Meier J, Sercu T, et al. Biological structure and function emerge from scaling unsupervised learning to 250 million protein sequences. Proc Natl Acad Sci USA. 2021;118(15):e2016239118. doi:10.1073/pnas.2016239118

Ryu JY, Kim HU, Lee SY. Deep learning enables high-quality and high-throughput prediction of enzyme commission numbers. Proc Natl Acad Sci USA. 2019;116(28):13996–14001. doi:10.1073/pnas.1821905116

Sanderson T, Bileschi ML, Belanger D, Colwell LJ. ProteInfer, deep neural networks for protein functional inference. eLife. 2023;12:e80942. doi:10.7554/eLife.80942

Steinegger M, Söding J. MMseqs2 enables sensitive protein sequence searching for the analysis of massive data sets. Nat Biotechnol. 2017;35:1026–1028. doi:10.1038/nbt.3988

UniProt Consortium. UniProt: the Universal Protein Knowledgebase in 2023. Nucleic Acids Res. 2023;51(D1):D523–D531. doi:10.1093/nar/gkac1052

Vaswani A, Shazeer N, Parmar N, et al. Attention is all you need. Adv Neural Inf Process Syst. 2017;30:5998–6008.

Yu T, Cui H, Li JC, Luo Y, Jiang G, Zhao H. Enzyme function prediction using contrastive learning. Science. 2023;379(6639):1358–1363. doi:10.1126/science.adf2465

