## Supplementary Table and Figure for "Protein Language Models Outperform BLAST for Evolutionarily Distant Enzymes: A Systematic Benchmark of EC Number Prediction"

#### Supplementary Figures

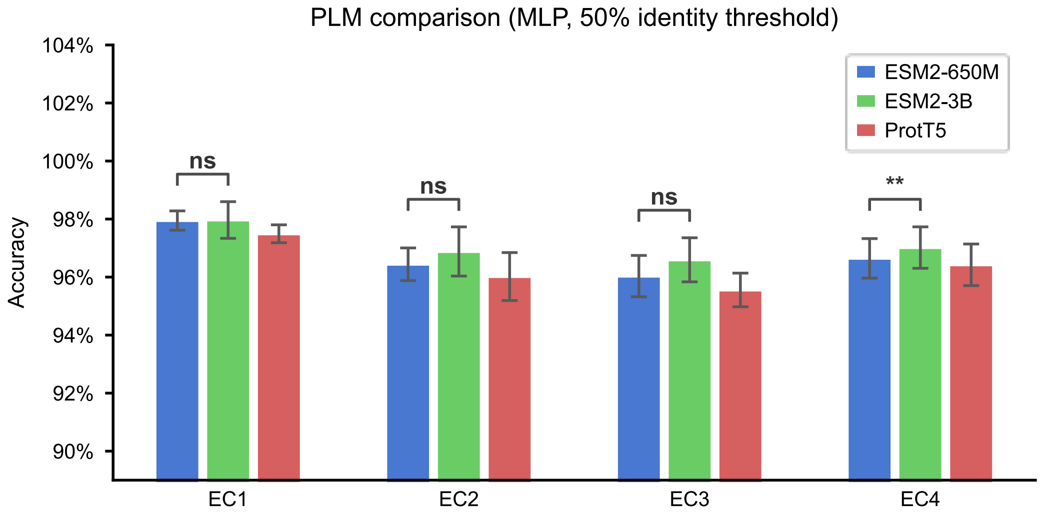


**Figure S1.** PLM comparison across EC levels. Grouped bar chart showing accuracy of ESM2-650M, ESM2-3B, and ProtT5-XL paired with MLP classifiers at 50% sequence identity threshold, for each EC hierarchy level. Error bars indicate ±1 standard deviation across three random seeds. Statistical comparison: ESM2-650M vs ESM2-3B ($p = 0.011$), ESM2-650M vs ProtT5 ($p = 0.003$), ESM2-3B vs ProtT5 ($p = 0.001$). Differences are statistically significant but small in absolute magnitude (≤0.35 pp between ESM2-650M and ESM2-3B).

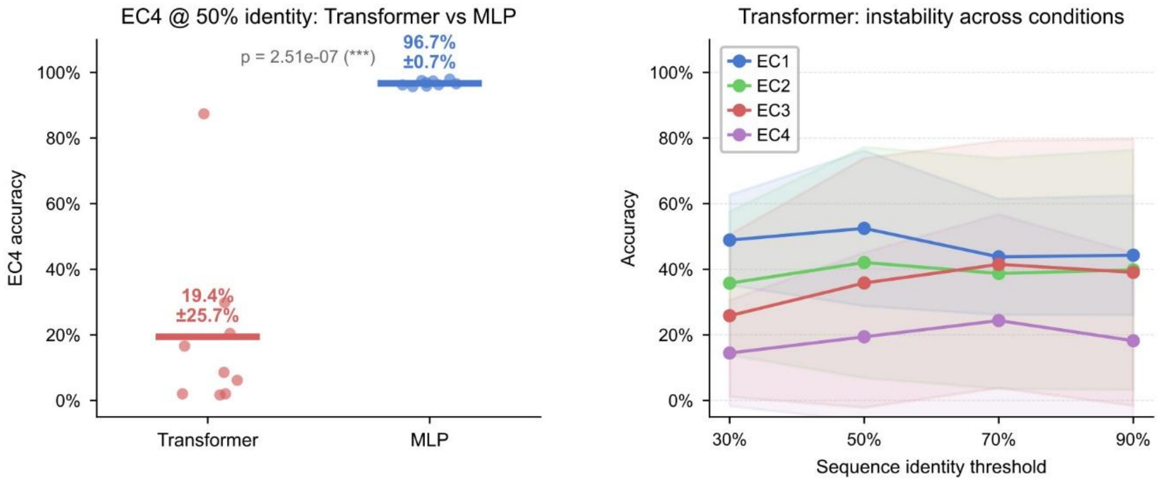


**Figure S2.** Transformer convergence instability. Left panel: strip plot comparing Transformer Encoder and MLP accuracy on EC4 at 50% sequence identity threshold. Each point represents one trained model (n = 9 per architecture: 3 PLMs × 3 seeds). The Transformer shows dramatically higher variance (45.9% ± 36.3%) compared to MLP (96.6% ± 0.8%). Right panel: Transformer accuracy as a function of sequence identity threshold for each EC hierarchy level. EC4 and EC1 both show extreme instability. This pattern is consistent with learning-rate-induced gradient instability (shared lr = $1\times10^{-3}$ is 10–100× too high for Transformer encoders).

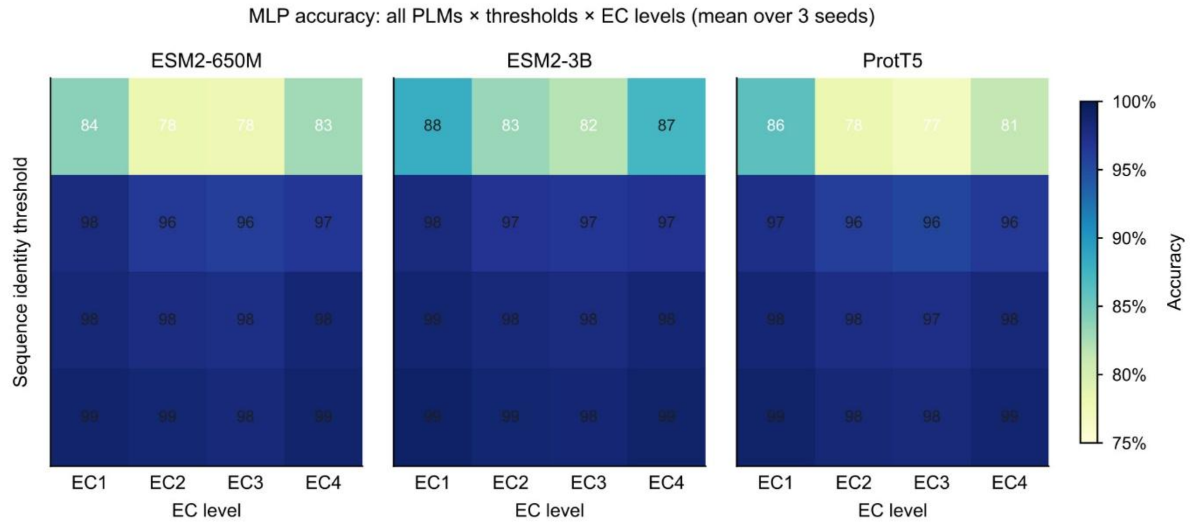


**Figure S3.** Full accuracy matrix: all PLMs × thresholds × EC levels. Three-panel heatmap showing mean accuracy (MLP classifier) for each PLM across all four sequence identity thresholds (rows: 30%, 50%, 70%, 90%) and EC hierarchy levels (columns: EC1–EC4). Values annotated in each cell. Color scale: YlGnBu, range 75%–100%. A single shared colorbar is placed to the right. All three PLMs show consistent threshold sensitivity; ESM2-3B is consistently highest, followed by ESM2-650M and ProtT5.

### Supplementary Tables

**Supplementary Table S1. Cross-organism validation, prokaryotes.**

| Organism | Domain | n (PLM) | PLM EC4 | BLAST EC4 | Δ |
|---|---|---|---|---|---|
| *Rhodopirellula baltica* | Bacteria | 2,735 | 88.9% | 54.9% | +34.0 pp |
| *Haloferax volcanii* | Archaea | 1,538 | 93.1% | 72.3% | +20.8 pp |
| *Sulfolobus acidocaldarius* | Archaea | 1,985 | 88.8% | 70.2% | +18.6 pp |
| *Methanococcus maripaludis* | Archaea | 1,255 | 91.4% | 75.5% | +15.9 pp |
| *Escherichia coli K-12 †* | Bacteria | 1,160 | 91.5% | 76.6% | +14.9 pp |
| *Deinococcus radiodurans* | Bacteria | 489 | 94.7% | 80.5% | +14.2 pp |
| *Prochlorococcus marinus MED4* | Bacteria | 1,828 | 89.2% | 76.3% | +12.9 pp |
| *Thermus thermophilus HB8* | Bacteria | 774 | 91.0% | 78.9% | +12.1 pp |
| *Buchnera aphidicola* | Bacteria | 1,663 | 93.8% | 85.4% | +8.4 pp |
| **Mean** | | | **91.4%** | **74.5%** | **+16.9 pp** |

ESM2-650M + MLP vs BLAST-90K, EC4, 50% identity threshold. Sorted by descending PLM−BLAST advantage. † E. coli K-12: ~20% of proteome in training set.

**Supplementary Table S2. Cross-organism validation, distant eukaryotes.**

| Organism | Category | n | PLM | BLAST-90K | BLAST-520K | Δ vs 90K |
|---|---|---|---|---|---|---|
| *Giardia lamblia* | Protist | 5,747 | 97.8% | 66.0% | 77.4% | +31.8 pp |
| *Trichomonas vaginalis* | Protist | 2,721 | 92.2% | 64.7% | 65.8% | +27.5 pp |
| *Leishmania major* | Protist | 1,226 | 88.9% | 78.2% | 81.6% | +10.7 pp |
| *Trypanosoma brucei* | Protist | 2,584 | 92.3% | 84.6% | 88.0% | +7.7 pp |
| *Cyanidioschyzon merolae* | Alga | 741 | 97.4% | 92.3% | 94.9% | +5.1 pp |
| *Mucor circinelloides* | Basal Fungus | 1,479 | 95.3% | 91.0% | 94.7% | +4.3 pp |
| *Encephalitozoon cuniculi* | Microsporidian | 353 | 96.0% | 93.8% | 95.2% | +2.3 pp |
| *Marchantia polymorpha* | Basal Plant | 5,272 | 95.0% | 93.6% | 97.0% | +1.4 pp |
| *Physcomitrella patens* | Basal Plant | 5,331 | 93.4% | 92.4% | 97.1% | +1.0 pp |
| *Rhizopus oryzae* | Basal Fungus | 3,180 | 91.8% | 91.0% | 94.7% | +0.8 pp |
| *Chlamydomonas reinhardtii* | Alga | 1,683 | 82.3% | 81.8% | 85.5% | +0.5 pp |
| *Plasmodium falciparum* | Protist | 9,037 | 76.5% | 87.2% | 89.3% | −10.7 pp |
| *Toxoplasma gondii* | Protist | 20,767 | 61.8% | 66.4% | 68.0% | −4.6 pp |

ESM2-650M + MLP vs BLAST, EC4, 50% identity threshold. Sorted by descending Δ vs BLAST-90K. Negative Δ = BLAST outperforms PLM.
